# Combination treatment can hinder the evolution of resistance to antimicrobial peptides

**DOI:** 10.1101/2022.03.13.484126

**Authors:** Bar Maron, Jonathan Friedman, Zvi Hayouka

## Abstract

Antibiotic resistant microbial pathogens are becoming a major threat to human health. Therefore, there is an urgent need to develop new alternatives to conventional antibiotics. One such promising alternative is antimicrobial peptides (AMPs), which are produced by virtually all organisms and typically inhibit bacteria via membrane disruption. However, previous studies demonstrated that bacteria can rapidly develop AMP resistance. Here, we study whether combination therapy, known to be able to inhibit the evolution of resistance to conventional antibiotics, can also hinder the evolution of AMP resistance. To do so, we evolved the opportunistic pathogen *S. aureus* in the presence of individual AMP, AMP pairs, and a combinatorial antimicrobial peptide library. Treatment with some AMP pair indeed hindered the evolution of resistance compared with individual AMPs. In particular, resistance to pairs was delayed when resistance to the individual AMPs came at a cost of impaired bacterial growth, and did not confer cross-resistance to other tested AMPs. The lowest level of resistance evolved during treatment with the combinatorial antimicrobial peptide library termed random antimicrobial peptide mixture, which contains more than a million different peptides. A better understanding of how AMP combinations affect the evolution of resistance is a crucial step in order to design ‘resistant proof’ AMPs cocktails that will offer a sustainable treatment option for antibiotic resistant pathogens.

## Introduction

Overuse of antibiotics in both medicine and agriculture has led to antibiotic resistant microorganisms becoming widespread in environmental and clinical settings. In particular, bacterial pathogens have become a major threat to public health, being responsible for more than 2.8 million infections and more than 35,000 deaths annually in the U.S alone [1]. *Staphylococcus aureus* is a significant human pathogen that causes multiple types of infections leading to morbidity and mortality [2,3]. It is known for its exceptional ability to develop resistance towards a multitude of antimicrobials [4].

Several approaches aiming at curbing the rise and spread of resistance have been proposed: (a) prudent use of antimicrobials; (b) development of new antimicrobials; and (c) development of treatment strategies that prevent or delay the evolution of antimicrobial resistance. Here we combine two of these approaches and study whether a treatment strategy based on combinations of antimicrobial peptides (AMPs) can hinder resistance development.

Antimicrobial peptides (AMPs) are a diverse family of compounds produced by virtually all organisms, [5–7] that typically inhibit bacteria via several mechanisms, mainly by disrupting bacterial membranes [8–12]. AMPs are considered to be a promising novel alternative to traditional antibiotics [13]. However, widespread application of AMPs may also cause the rise of AMP resistant pathogens. Rapid evolution of resistance to AMPs will negate their efficacy and may compromise the activity of AMPs that are part of the immune system. Therefore, there is a need to develop treatment strategies involving AMPs that delay the evolution of resistance to these antimicrobial agents.

Recent studies have shown that resistance to AMPs can evolve in the lab and in nature [14,15]. Several *in vitro* studies showed that AMPs can select for resistant bacteria [16]. Perron *et al*. evolved *Escherichia coli* and *Pseudomonas fluorescens* in the presence of pexiganan – a synthetic analogue of frog antimicrobial peptides (magainins) [17]. Both bacterial species evolved resistance to pexiganan within approximately 650 generations. In addition, Dobson *et al*. evolved gram-positive bacteria *Staphylococcus aureus* in the presence of different antimicrobials including antimicrobial peptides and showed that *S. aureus* evolved resistance against AMPs, albeit slower than against antibiotics [18]. A slower rate of evolution was also observed against the prokaryotic AMP colistin, when compared with antibiotics [19].

Treatment with a combination of antimicrobials may hinder the rise of resistance [20], but only few studies have been investigating the evolution of resistance towards AMPs combinations [18,21]. Dobson *et al*. found that *S. aureus* went extinct more rapidly when treated with a mixture of two antimicrobial peptides – pexiganan and melittin - compared to single AMP treatments [18]. However, it is still not clear how general are these results, and which AMP combinations will lead to delayed resistance evolution.

In this study, we aim to elucidate the evolution of resistance towards individual AMPs and their combinations, and the factors that influence the combination’s efficiency. To do so, we have performed experimental evolution of *S. aureus* in the presence of several individual AMPs and AMP combinations. A better understanding of how AMP combinations affect the evolution of resistance is a crucial step in order to the development of ‘resistant proof’ AMPs cocktail that can offer a sustainable treatment option of antibiotic resistant pathogens [22].

## Results

### Resistance towards single AMPs evolved readily

First, we performed experimental evolution of *S. aureus* in the presence of six individual AMPs with different modes of bacterial membrane distruption (Table S1). Prior to the experimental evolution the antimicrobial activity of each AMP towards *S. aureus* was evaluated by measuring its minimal inhibition concentration (MIC) (Table S1). The evolution experiment was designed to maintain strong selection for resistance, yet to avoid extinction. Therefore, each of six replicate lines of bacteria was exposed to four concentrations of each AMP: 1.5xMIC, 1xMIC, 0.5xMIC, 0.25xMIC (Fig 1A). Every four transfers the MIC was doubled if four out of six lines grew in the MIC or higher. Before the selection, we inoculated the bacteria with AMP free medium (MHB) in order to habituate it to the experimental conditions. The bacteria were transferred each day to fresh media (diluted 1:20) for seven days. The first experimental evolution assay was performed with six individual AMPs for 29 days (≈ 130 generations), each AMP had six parallel independent lines of bacteria.

**Fig 1.**
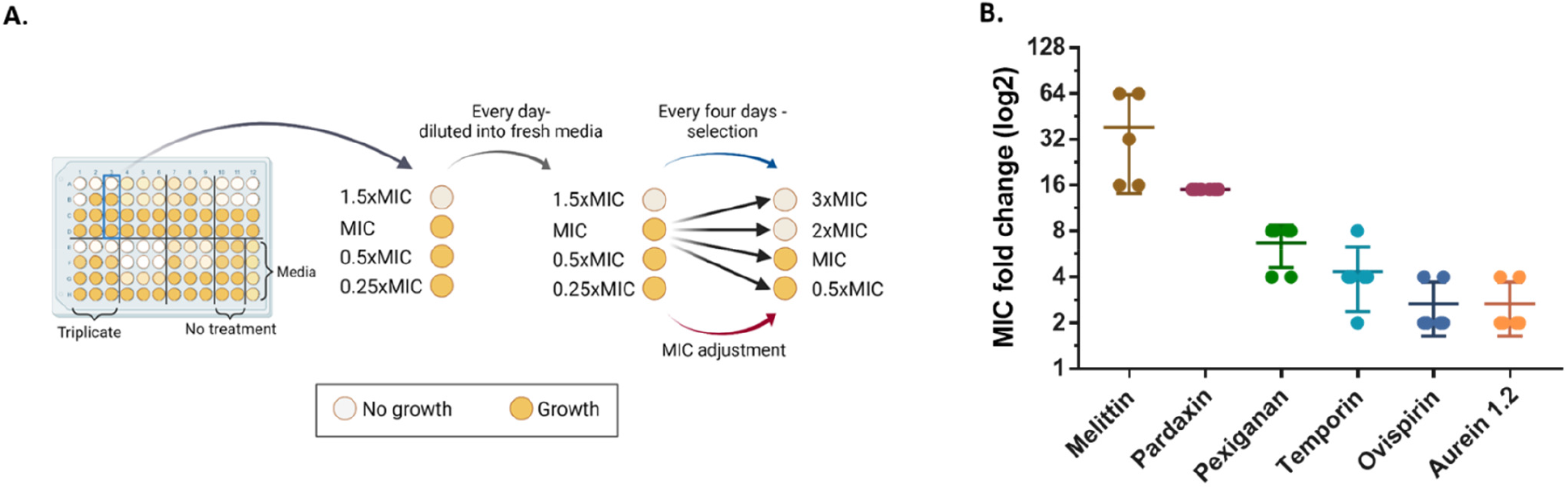
Resistance evolution varied between AMPs. (A) Experimental evolution procedure. Every day, 10μl microliters of last day well copied into new plate with fresh media and AMP/antibiotics. Every 4 days, bacteria that grew in highest AMP concentration selected and transferred into the four new concentrations of the line (black arrows). AMPs/antibiotics concentrations doubled when growth observed in 4 out of 6 lines in MIC concentration or above. Each 96-well plate contained: triplicate of each treatment (in two separate plates, total of 6 lines), 8 wells of bacteria without treatment, and media control. The experiment continued for approximately 130 generations 29 days). Evolved strains were isolated after 29 days. (B) Resistance of evolved strains. Resistance determined by MIC assay of each strain towards the corresponding AMP. Results shown as fold change of ancestor MIC values (n=6, n_mel_ =5, bars represent mean ±SD).

At the end of the experiment, the evolved strains were isolated, and the MIC values were determined in order to evaluate the level of resistance in the same conditions for all strains. The level of resistance that evolved varied significantly across AMPs (Fig 1B, p<0.0001). The strains that evolved with melittin had the largest increase in resistance, with mean MIC which was 38-fold higher than the ancestor strain. The MIC of pexiganan- and temporin-evolved strains increased by a 5- and 4-fold (respectively). Interestingly, ovispirin and aurein for which the MIC increased the least (2-fold on average) both have a carpet model mechanism of action [23,24] as oppose to the other AMPs.

### Combinations of AMPs can hinder the evolution of resistance

To further investigate the evolution of resistance with AMPs combinations, we performed experimental evolution with the AMPs for which medium-high resistance evolved: melittin, pexiganan and temporin (pardaxin AMP was excluded due to solubility issues). Before we started the evolution experiment with AMPs combinations, we evaluated the interaction between AMPs using checkboard assay. We found that there is no notable synergistic/antagonistic interaction between the tested combinations (Table S2). Therefore, the initial MIC concentration of each combination contained 0.5xMIC of each AMP. The experimental evolution procedure was performed similar to the first experiment (approximately 130 generations).

Resistance evolution towards individual AMPs in this experiment was consistent with the results of the first experiment (Fig 2). Temporin-evolved strains exhibit medium-low resistance, melittin-evolved strains had relatively high resistance. For the combination treatments, temporin-melittin could not inhibit the evolution of resistance better than temporin alone (Fig 2A). However, the combination of temporin-pexiganan hindered the evolution of resistance compared with each AMP alone (Fig 2B). The combination of pexiganan-melittin showed an even stronger ‘synergistic’ effect, where the evolved bacteria managed growing in an AMP concentration only 4-fold than the initial MIC, whereas they could grow in concentration 10-20-fold higher the initial MIC when evolved with the individual AMPs (Fig 2C, S1). Overall, treatment with some AMP combinations resulted in drastically lower resistance than treatment with each individual compound.

**Fig 2.**
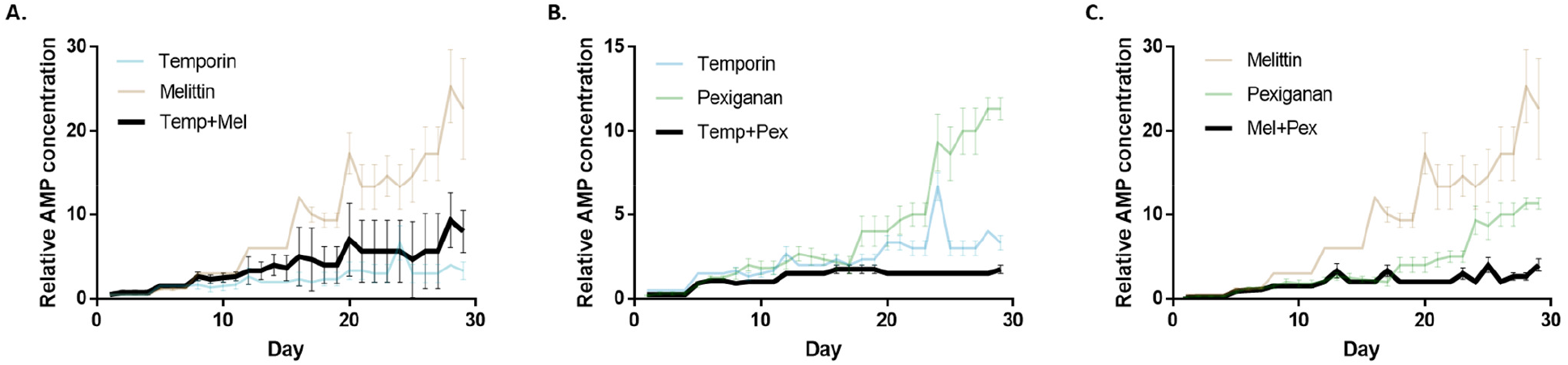
AMP combinations can hinder the evolution of resistance. Data represent the concentration of AMPs where bacteria grew, normalized to initial MIC (n=6, mean ± SE, Growth defined as OD_595nm_>0.3). (A) temporin-melittin combination. (B) temporin-pexiganan combination. (C) melittin-pexiganan combination. Each graph shows the concentrations of individual AMPs and their combinations through the evolution. Data of individual AMPs is identical in all panels. Mel, Melittin; Pex, Pexiganan; Temp, Temporin.

To further verify our findings, we isolated strains from the last day of the experiment and performed a standard MIC assay. The results of this MIC assay are consistent with the resistance levels found during the experimental evolution (Fig 3). All the combinations that contained melittin were found to reduce the resistance towards it. Pexiganan combinations result in the most effective combinations, as the MIC value towards pexiganan drops from an average of 9-fold change to 3-fold change in the combinations. The results from temporin-evolved strains confirm that these combinations were less effective compared to temporin alone, except for the pexiganan-temporin combination.

**Fig 3.**
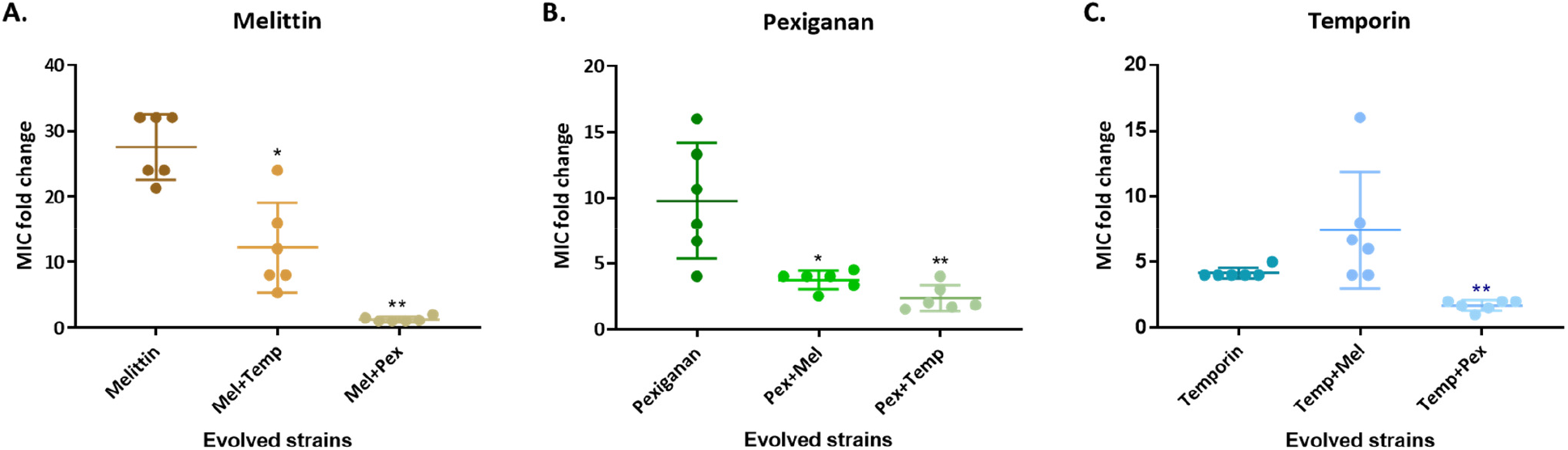
Resistance towards melittin and pexiganan was significantly reduced in most combinations. Each strain isolated from the end of the experiment and MIC values were determine using a standard assay. (A) melittin-evolved combinations, (B) pexiganan-evolved combinations, (C) temporin-evolved combinations. The MIC values refer to the AMP in the title, and normalized to ancestor’s MIC. Each dot represents a line (n=6), bars represent mean ± SD. * p< 0.05, ** p< 0.01 (Mann-Whitney U test with Bonferroni correction. Comparison of each combination to the individual-AMP’s strain). Mel, Melittin; Pex, Pexiganan; Temp, Temporin. The results of each dot represent the mean of at least three independent experiments.

### Resistance to melittin and pexiganan incurred a notable fitness cost

Next, we wanted to understand why some combinations are more effective than other in inhibiting the resistance occurrence. One hypothesis is that if there is a significant fitness cost of resistance to individual AMPs, the combination of two high-cost AMPs will better hinder the evolution of resistance. To test this hypothesis, we measured the growth ability of each evolved strain in the absence of AMPs (Fig 4).

**Fig 4.**
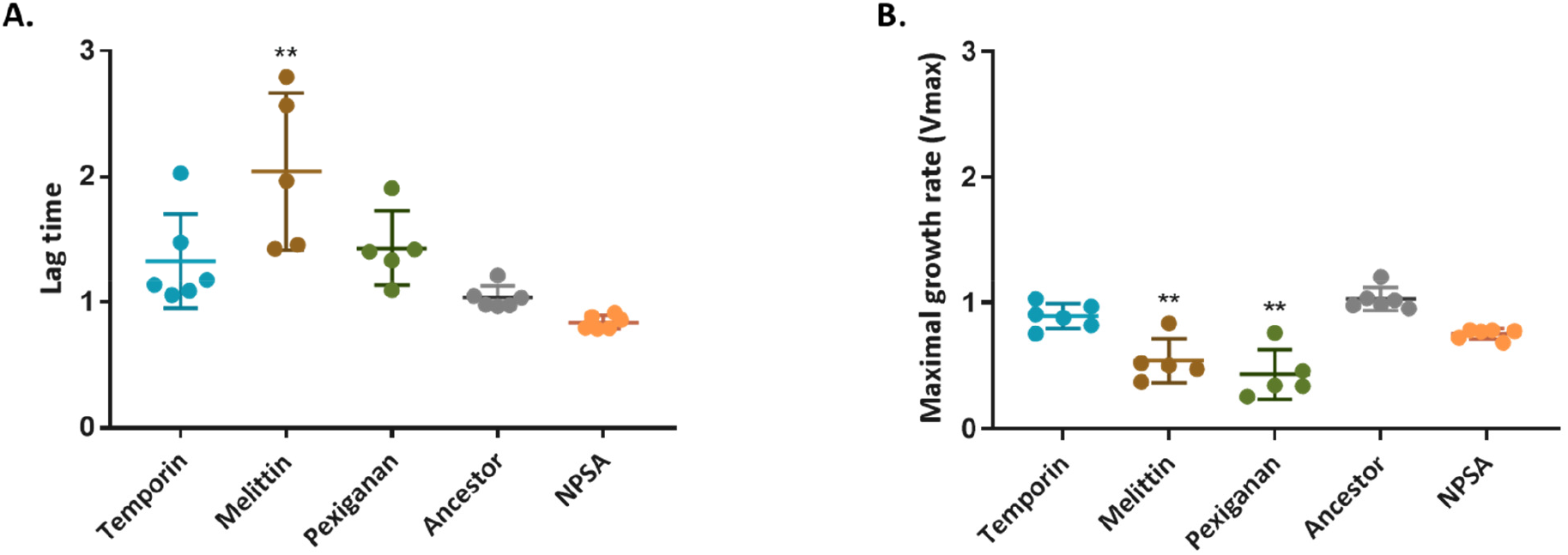
Resistance to melittin and pexiganan incurs a fitness cost in the form of a longer lag time and reduced growth rate. Determination of fitness cost performed by growing the bacteria in the absence of AMPs. OD_595nm_ was measured every 15 min through 24 hours. (A) lag time. (B) maximal growth rate (V_max_). Values were calculated by plate reader software (Gen 5 and normalized to ancestor strain values). Each dot represents mean of triplicate, bars represent mean ± SD (n_melittin, pexiganan_=5, Dunn’s multiple comparison refed to ancestor strain, ** p< 0.01). NPSA-evolved strain without AMPs. The results represent three independent experiments.

The lag time increased significantly in melittin strains (Fig 4A, p=0.0078), and a similar increase was found in the time to achieve maximal growth rate (Fig S2). Lag time also increased in temporin and pexiganan strains but not statistically significantly (p_Temp_=0.6585, p_Pex_=0.3323). Another important aspect of bacterial fitness is the maximal growth rate, which was reduced in both pexiganan and melittin strains, but not in temporin-evolved strains (Fig 4B). The overall fitness, which can be expressed by the area under the growth curve was significantly attenuated in pexiganan-evolved strains (p=0.0095) and melittin-evolved strains (p=0.05) (Fig S2). The fitness cost of the strains that evolved with AMP combinations was not impaired, except for temporin-melittin maximal growth rate (Fig S3). Overall, these results indicate that the AMPs for which resistance incurred the largest fitness costs, melittin and pexiganan, are indeed the ones whose combination most considerably hindered the evolution of resistance.

### Cross-resistance evolved but not towards all AMPs

Another possible mechanism affecting the evolution of resistance to combination therapy is cross-resistance or collateral sensitivity. If resistance towards a single AMP confers resistance to another AMP (cross-resistance), we hypothesize that a combination of these AMPS will be less effective at delaying the evolution of resistance. However, when resistance confers collateral sensitivity towards another AMP, we expect that this combination will be more effective. To examine this hypothesis, we determined the MIC value of each of the evolved strains towards the other AMPs it has not evolved with (Fig 5). We found that cross-resistance towards temporin was common (Fig 5A), which may explain why it was not effective at delaying resistance in combination treatments. Cross-resistance to pexiganan was rare, and temporin-evolved strains even showed increased sensitivity to pexiganan. These results are consistent with the fact that AMP combinations that contain pexiganan were more effective for delaying resistance evolution. Notably, melittin- and pexiganan-evolved strains were the only AMP pair where no major cross-resistance occurred, which may be another factor contributing to it being the combinations for which resistance was most drastically reduced.

**Fig 5.**
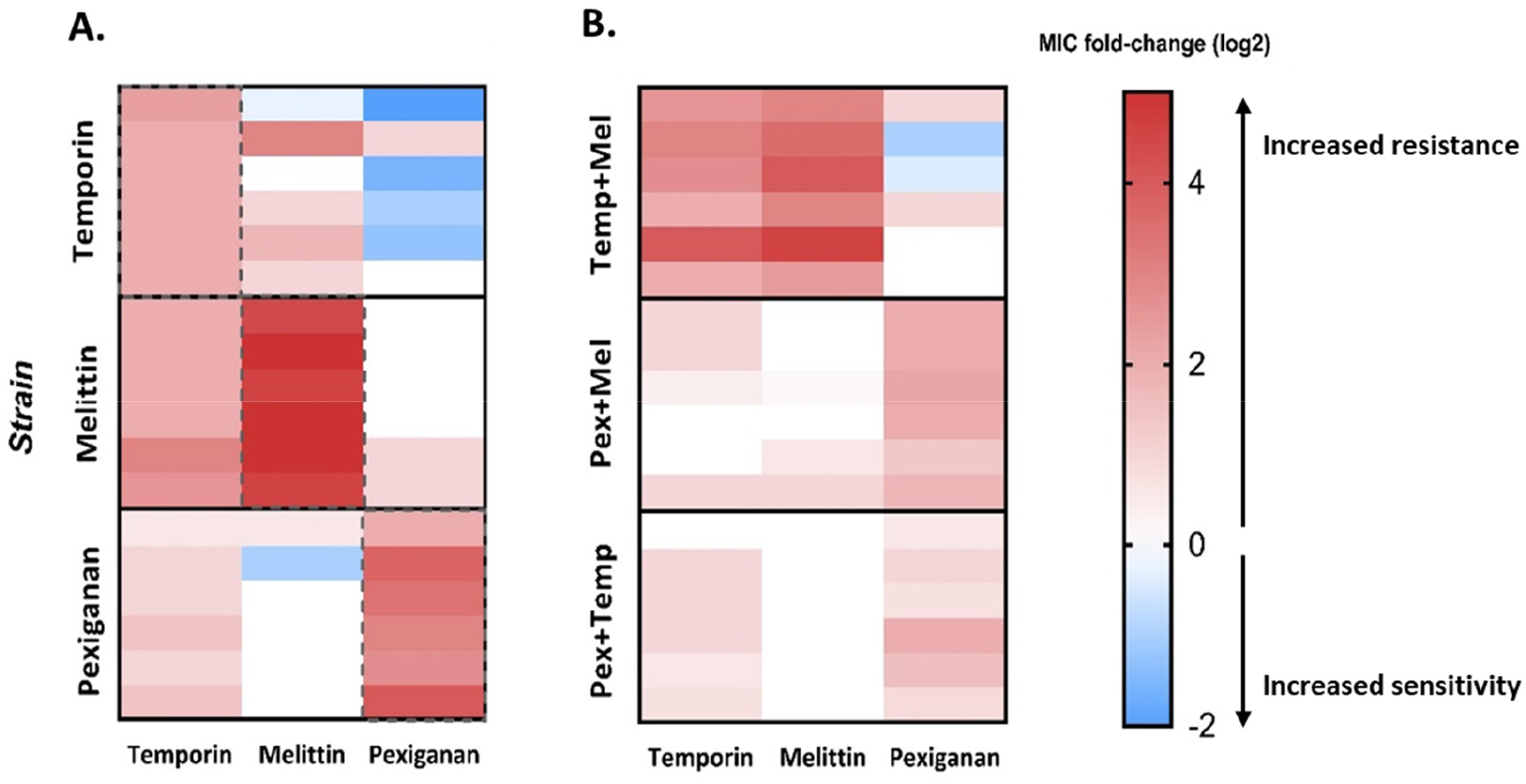
Cross-resistance evolved frequently towards temporin but less towards pexiganan. A standard minimal-inhibitory concentration assay (MIC) was performed in order to evaluate the cross-resistance and collateral sensitivity. In each experiment, the evolved strain was exposed to different AMPs, and the selected AMP as control. The MIC values present as fold change (log2) of the ancestor’s MIC. The results represent the mean of three independent experiments. (A) individual AMP evolved strains. (B) Two-AMPs evolved strains. Red color indicates resistance, blue indicates sensitivity. Dashed lines indicate the MIC towards the AMP it evolved with (not cross-resistance or sensitivity).

To further explore our hypothesis, we have also assayed the resistance of strains evolved with AMP combinations to all three individual AMPs (Fig 5B). Strains that evolved with melittin-temporin combination exhibited medium resistance, corresponding to the individual AMPs in Figure 5A. The strains that evolved in combination with pexiganan presented resistance towards pexiganan but less towards melittin, even when evolved with melittin-pexignan combination. Overall, except for three strains out of eighteen, bacteria remained susceptible to at least one AMP, which suggests that resistance towards some AMPs does not confer a general cross-resistance towards another AMPs.

### *S. aureus* developed low resistance towards a combinatorial antimicrobial peptide library

Given that pairs of AMPs hindered resistance evolution, we next tested whether more diverse AMP combinations may delay resistance evolution even further. We explored a novel type of AMPs: random antimicrobial peptides mixture, a combinatorial library of antimicrobial peptides (RPMs) [25]. The peptides in the mixture contain only two type of amino acids, one hydrophobic and one cationic, and defined chain length of 20 amino acids. Thus, the RPM contains more than a million sequences (2^20^ optional sequences in the mixture) of peptides that are all composed of the same two amino acids. RPMs were found to be highly effective against a variety of pathogens, even *in vivo* [26]. We evolved the bacteria with 20-mer RPM composed of phenylalanine and lysine (FK), which represents a complexed case of peptides combinations. The MIC to this RPM increased only by a factor of two throughout the entire course of our experimental evolution (Fig. 6A), a lower increase than for any of the individual AMP and 2-AMPs combinations (Fig S4).

**Fig 6.**
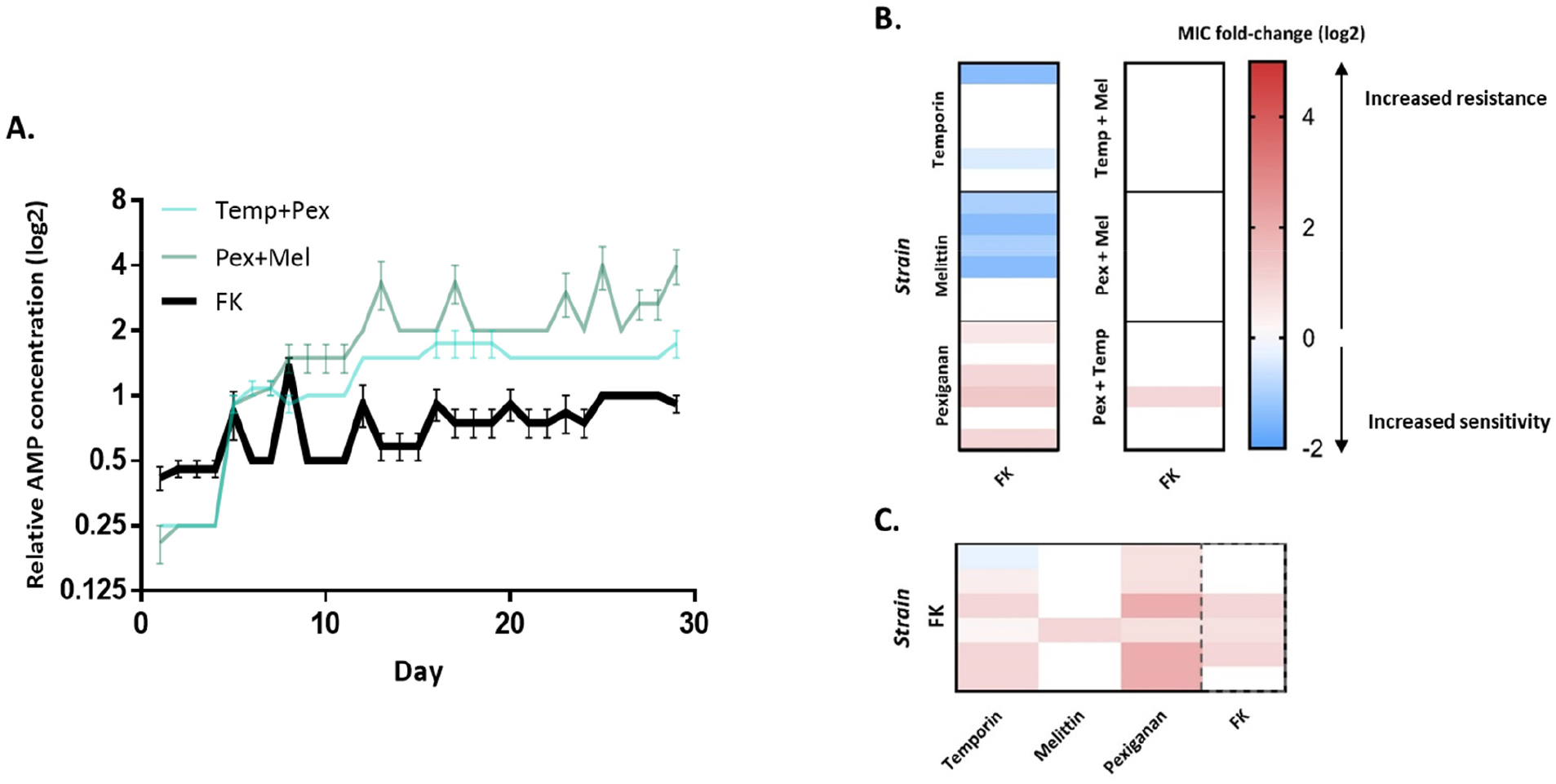
Random-peptides mixture (FK) hinder the evolution of resistance and cross-resistance towards it. (A) Data represent the concentration of AMPs where bacteria grew, normalized to initial MIC (n=6, mean ± SEM, Growth defined as OD_595nm_ >0.3. Each graph shows the concentrations of AMPs through the evolution. FK-random-peptides mixture. (B) Susceptibility of evolved strains towards FK20. (C) Susceptibility of FK evolved strains towards different AMPs. Data in B and C represent the mean of three independent experiments. Dashed lines indicate the MIC towards the AMP it evolved with (not cross-resistance or sensitivity).

We further investigated the sensitivity of the strains that evolved with individual AMP to the RPM (FK). Surprisingly, six out of 18 strains develop collateral sensitivity, and nine strains maintained a similar sensitivity (Fig 6B). Cross-resistance to FK RPM occurred in four out of six pexiganan-evolved strains. Except for pexiganan strains, no cross-resistance evolved towards FK (only 1/30). Low cross-resistance to individual AMPs also occurred in most of FK-evolved strains (Fig 6C), though, the strains remain sensitive to melittin (except of one strain). Yet, three out of the six strains remained sensitive as the ancestor (towards FK), as the rest evolved resistance that increase the MIC by 2-fold maximum. In summary, these results indicate that the AMP-resistant strains could be treated using AMP combinations, with the same efficacy as the ancestor’s. Furthermore, development of general resistance towards all AMPs is less likely to occur.

## Discussion

Antimicrobial peptides are naturally found in most organisms, as part of their immune system. Typically, AMPs are found in diverse mixtures of different peptides or other type of antimicrobial agents rather than individual ones [27–32]. Yet, there are only a few studies that investigated the evolution of bacteria in the presence of more than one AMP. In the current study, we examined the evolution of resistance of the bacterial pathogen *S. aureus* towards AMP pairs, and the factors that impact the rate with which resistance arises.

We first demonstrated that *S. aureus* can readily evolve resistance towards selected individual AMPs. The resistance that was observed varied between 2 to 64-fold increase in the MIC values. Interestingly, the lowest resistance evolved towards AMPs that act on the bacterial membrane via carpet model: ovispirin and aurein. In contrast, the highest resistance evolved against pore-forming AMPs (pexiganan, melittin, pardaxin), with temporin being an exception. Further, we have observed that a combination of two AMPs can delay the evolution of resistance, compared to each individual AMP. These results are consistent with Dobson *et al*., who performed experimental evolution with a combination of pexiganan and melittin.

To further explore the reason for the low resistance occurrence we quantified the fitness cost effect. We have shown that the fitness cost of resistance to AMPs differs by the strains, and that melittin and pexiganan strains had the most impaired growth fitness. The combinations of these two AMPs substantiallyhindered the evolution of resistance compared to each AMP alone, suggesting that the high cost of resistance to these AMPs can be an explanation for this combination’s effective reduction of resistance evolution. Antibiotic-resistance strains that are clinically isolated are frequently found to have similar fitness in the absence of antibiotics, which strengthen the hypothesis that combinations of high-cost resistance AMPs might be effective for preventing resistance occurrence [33].

Moreover, temporin strains evolved collateral sensitivity towards the AMP pexiganan. This result may contribute to the effectiveness of the combination of temporin-pexiganan. The fact that temporin-resistant strains did not show a fitness cost could select for these strains rather than pexiganan-resistant strains, and therefore become more sensitive to pexiganan. In contrast, temporin and melittin strains both showed cross-resistance towards each other, which might be a reason for this combination to be less efficient. These results are consistent with previous studies that showed that the collateral-sensitivity between antibiotics can limit resistance evolution [35,36]. However, Nichol *et al*., suggested that collateral sensitivity is never universal, and that cross-resistance could be developed if a different evolutionary trajectory is taken [37].

Cross-resistance between AMP-resistant strains is raising concerns since many of them have a shared mechanism of action – disruption of the bacterial membrane [34]. We found that although our selected AMPs act on the membrane, they are not sharing cross-resistance across all AMPs. In addition, AMP-resistance strains were still sensitive to a combination of AMP (FK). These results suggest that cross-resistance to immune system AMPs may not evolve so readily in due to clinical AMP treatments.

Treatment with a combinatorial library of over a million AMP sequences resulted in the lower levels of resistance than treatments involving AMP pairs. This suggests that cocktails involving multiple AMP may be more effective in hindering the evolution of resistance. Nevertheless, it is still unclear whether this requires ultra-diverse cocktails, such as the random AMP library include in this study, or whether a defined mixture of several AMPs may suffice. Overall, these results emphasize the potential of AMPs combinations to hinder resistance evolution and support the rational of using antimicrobial peptides mixture rather than individual AMPs.

In summary, we have demonstrated the ability of antimicrobial peptide combinations to delay the evolution of resistance in the pathogen *S. aureus*. Further research is needed in order to uncover the genetic and mechanistic basis of resistance to AMPs and to test to what extent the trends we found hold more broadly across other antimicrobials and pathogens.

## Materials and methods

### Bacterial strains and growth conditions

All experiments were performed with *S. aureus* JLA513 [38] (*hla-lacZ hla*+, derived from SH1000, kindly provided by Jens Rolff) which contains tetracycline resistance. This strain is defined as the ancestor strain. Prior to each experiment, strains were isolated from Muller-Hinton (MH, HiMedia) ager plates and individual colonies were picked and grown in MH broth overnight in 37 °C. All bacterial cells used in this study were stored in 25% glycerol at -80 °C.

### Synthesis of antimicrobial peptides

Six different AMPs with three different modes of action were selected for the experimental evolution assay (see Table S1). All AMPs were synthesized using Fmoc solid-phase peptide synthesis method (SPPS) using Peptide synthesizer (Liberty Blue, CEM, USA). Upon synthesis completion, peptides were cleaved from the resin [(95 % trifluoroacetic acid (TFA), 2.5 % water, 2.5 % triisopropylsilane (TIPS)], re-suspended in double distilled water (DDW), frozen and lyophilized. Subsequently, the crude peptide dissolved in DMSO and purified using semi-preparative RP-HPLC, while MALDI-TOF-MS utilized for verification of the peptides mass and purity. Random peptide mixtures (RPM) containing phenylalanine and lysine (FK20) were synthesized as previously described [39].

### Minimal-inhibitory concentration (MIC) determination

MIC values were determined using a standard protocol [40]. Briefly, *S. aureus* cells were grown overnight in MH at 37 °C, 200 rpm. Subsequently, cells were diluted 1:100 in MH and grown until reaching OD_595nm_ 0.1. Then, 100μl of 5×10^5^ CFU/ml were inoculated into each well in 96-well plates that contained a serial dilution of AMPs. Each plate contained 3 replicates of each AMP. MIC value were defined as the lowest concentration at which there is inhibition of bacterial growth by at least 90%. Fold change of MIC in evolved strains was divided by the ancestor MIC.

Cross-resistance and collateral sensitivity were evaluated by the same method as MIC assay. Each strain was exposed to different AMPs, and the AMP that it evolved with as control. Ancestor’s MICs were determined as well in each experiment. Each experiment was repeated at least twice independently.

### Experimental evolution procedure

Prior to evolution with AMPs, an *S. aureus* JLA513 colony was transferred from MH agar plate into 5 ml MH broth in a 50 ml tube, and incubated overnight in 37 °c, 200 rpm. Subsequently, this starter culture was diluted by 20-fold into a 1.5 ml Eppendorf containing 850 μl MH broth to maintain the same headspace ratio as in the experimental evolution procedure and incubated in the same conditions (37 °c, 200 rpm) overnight. These dilutions by 20-fold were repeated for 7 transfers in order to habituate the bacteria to the experimental conditions. Our experimental evolution procedure was designed to exert selective pressure, yet to avoid extinction of lines. Therefore, each line was exposed to 4 concentrations of AMP according to its MIC: 1.5x, 1x, 0.5x, 0.25x (Figure 1, Table S1). Experiments were performed in 96-well plates and each AMP had 6 parallel lines (same ancestor). In the AMPs combination treatments, the effective ratio between the AMPs was 1:1, as the MIC concentration contained 0.5xMIC of each. In each plate, 8 wells with bacteria only (no AMPs) were used as a positive control. Four wells with media only serve as negative control, to indicate contaminations. Six lines that evolved with rifampicin were used as positive control for resistance evolution, as *S. aureus* evolves resistance towards it readily [41]. Every day, 10 μl of the previous plate were replicated into 190 μl of fresh media and AMPs. Every 4 days, bacteria from the highest concentration of AMP were selected and transferred into 4 concentrations in the new plate. MIC was doubled when growth was observed in 4 out of 6 lines, in MIC concentration or higher. Growth was defined as O.D>0.3. The experimental evolution was carried out for 29 transfers, which are approximately 122 generations. Before every selection or MIC increment, samples were taken to make glycerol stocks (25%) and preserved in -80°c, to avoid lines extinction. Spot plating was performed on MHA contains 5 μg/ml tetracycline to indicate growth before selection.

### Fitness cost of evolved strains

Bacterial cells were grown overnight in MH, then diluted to OD_595_ 0.1/100. Two hundred microliters of each strain’s culture were transferred into 96-well plates. Each strain had 3 repeats in the same plate. Optical density (OD_595_) was measured every 15 min through 24 h using Epoch 2 microplate reader (BioTek). Lag time, Vmax and t at Vmax were calculated using the plate reader software (Gen5) and normalized to the ancestor strain’s values. The area under the curve was calculated by OD_595_ measurements until stationary phase (14 h).

### AMPs interaction assessment using checkerboard assay

Before proceeding to evolution with combination of AMPs, the interaction between AMPs has been assessed. In order to evaluate these interactions, checkerboard assays were performed as previously described [40]. Briefly, hundred microliters of 5×10^5^ CFU/ml log phase *S. aureus* cells were added into 96-well plates with different concentrations of two AMPs. The plates were then incubated for 24 h in 37°c and bacterial growth was determined by measuring optical density (OD_595nm_) using plate reader. The checkerboard assay results were used to calculate the Fractional Inhibitory Concertation index (FICi) according to the following equation:

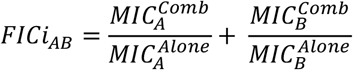

Where 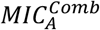 is the MIC of peptide A when combined with peptide B, and 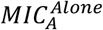 is the MIC peptide A alone, without the presence of compound B. Finally, the overall interactions between peptides were quantified in two methods: minimal FIC and average FIC. The minimal FIC is the lowest concentration (FIC) where bacteria were inhibited by at least 80%. In the average FIC method, we calculated the FIC for each concentration that bacteria were inhibited by at least 80%, and then the average FIC for all values was calculated. Each experiment was performed in duplicate. The results represent two independent experiments.

## Notes

### Competing Interest Statement

The authors have declared no competing interest.

## References

1. Nelson RE, Hatfield KM, Wolford H, Samore MH, Scott II RD, Reddy SC, et al. National Estimates of Healthcare Costs Associated With Multidrug-Resistant Bacterial Infections Among Hospitalized Patients in the United States. Clin Infect Dis. 2021;72: S17–S26. doi:10.1093/cid/ciaa1581

2. Lee GC, Dallas SD, Wang Y, Olsen RJ, Lawson KA, Wilson J, et al. Emerging multidrug resistance in community-associated Staphylococcus aureus involved in skin and soft tissue infections and nasal colonization. J Antimicrob Chemother. 2017;72: 2461–2468. doi:10.1093/jac/dkx200

3. Loir Y Le, Baron F, Gautier M, Loir Y Le, Baron F, Gautier M, et al. Staphylococcus aureus and food poisoning. Genet Mol Res. 2003;2: 63–76.

4. Foster TJ. Antibiotic resistance in Staphylococcus aureus. Current status and future prospects. FEMS Microbiol Rev. 2017;41: 430–449. doi:10.1093/femsre/fux007

5. Diamond G, Beckloff N, Weinberg A, Kisich KO. The Roles of Antimicrobial Peptides in Innate Host Defense. Curr Pharm Des. 2009;15: 2377–92. doi:https://doi.org/10.2174/138161209788682325

6. Michael Zasloff. Antimicrobial peptides of multicellularorganisms. Nature. 2002;415: 389–395. doi:https://doi.org/10.1038/415389a

7. Rathinakumar R, Walkenhorst WF, Wimley WC. Broad-spectrum antimicrobial peptides by rational combinatorial design and high-throughput screening: The importance of interfacial activity. J Am Chem Soc. 2009;131: 7609–7617. doi:10.1021/ja8093247

8. Zasloff M, Martin B, Chen HC. Antimicrobial activity of synthetic magainin peptides and several analogues. Proc Natl Acad Sci U S A. 1988;85: 910–913. doi:10.1073/pnas.85.3.910

9. Zasloff M. Antimicrobial peptides, innate immunity, and the normally sterile urinary tract. J Am Soc Nephrol. 2007;18: 2810–2816. doi:10.1681/ASN.2007050611

10. Rathinakumar R, Wimley WC. Biomolecular engineering by combinatorial design and high-throughput screening: Small, soluble peptides that permeabilize membranes. J Am Chem Soc. 2008;130: 9849–9858. doi:10.1021/ja8017863

11. Oren Z, Shai Y. Selective lysis of bacteria but not mammalian cells by diastereomers of melittin: Structure-function study. Biochemistry. 1997;36: 1826–1835. doi:10.1021/bi962507l

12. Oren Z, Ramesh J, Avrahami D, Suryaprakash N, Shai Y, Jelinek R. Structures and mode of membrane interaction of a short α helical lytic peptide and its diastereomer determined by NMR, FTIR, and fluorescence spectroscopy. Eur J Biochem. 2002;269: 3869–3880. doi:10.1046/j.1432-1033.2002.03080.x

13. Aoki W, Ueda M. Characterization of antimicrobial peptides toward the development of novel antibiotics. Pharmaceuticals. 2013;6: 1055–1081. doi:10.3390/ph6081055

14. Spohn R, Daruka L, Lázár V, Martins A, Vidovics F, Grézal G, et al. Integrated evolutionary analysis reveals antimicrobial peptides with limited resistance. Nat Commun. 2019;10. doi:10.1038/s41467-019-12364-6

15. Andersson DI, Hughes D. Mechanisms and consequences of bacterial resistance to antimicrobial peptides. Drug Resist Updat. 2016;26: 43–57. doi:10.1016/j.drup.2016.04.002

16. Kubicek-Sutherland JZ, Lofton H, Vestergaard M, Hjort K, Ingmer H, Andersson DI. Antimicrobial peptide exposure selects for Staphylococcus aureus resistance to human defence peptides. J Antimicrob Chemother. 2017;72: 115–127. doi:10.1093/jac/dkw381

17. Perron GG, Zasloff M, Bell G. Experimental evolution of resistance to an antimicrobial peptide. 2006; 251–256. doi:10.1098/rspb.2005.3301

18. Dobson AJ, Purves J, Kamysz W, Rolff J. Comparing Selection on S. aureus between Antimicrobial Peptides and Common Antibiotics. PLoS One. 2013;8. doi:10.1371/journal.pone.0076521

19. Suzuki S, Horinouchi T, Furusawa C. Prediction of antibiotic resistance by gene expression profiles. Nat Commun. 2014;5. doi:10.1038/ncomms6792

20. Dean Z, Maltas J, Wood KB. Antibiotic interactions shape short-term evolution of resistance in E. Faecalis. PLoS Pathog. 2020;16: 1–24. doi:10.1371/journal.ppat.1008278

21. El Shazely B, Yu G, Johnston PR, Rolff J. Resistance Evolution Against Antimicrobial Peptides in Staphylococcus aureus Alters Pharmacodynamics Beyond the MIC. Front Microbiol. 2020;11: 1–11. doi:10.3389/fmicb.2020.00103

22. Bell G, MacLean C. The Search for ‘Evolution-Proof’ Antibiotics. Trends Microbiol. 2018;26: 471–483. doi:10.1016/j.tim.2017.11.005

23. Yamaguchi S, Huster D, Waring A, Lehrer RI, Kearney W, Tack BF, et al. Orientation and dynamics of an antimicrobial peptide in the lipid bilayer by solid-state NMR spectroscopy. Biophys J. 2001;81: 2203–2214. doi:10.1016/S0006-3495(01)75868-7

24. Giacometti A, Cirioni O, Riva A, Kamysz W, Silvestri C, Nadolski P, et al. In vitro activity of aurein 1.2 alone and in combination with antibiotics against gram-positive nosocomial cocci. Antimicrob Agents Chemother. 2007;51: 1494–1496. doi:10.1128/AAC.00666-06

25. Hayouka Z, Bella A, Stern T, Ray S, Jiang H, Grovenor CRM, et al. Binary Encoding of Random Peptide Sequences for Selective and Differential Antimicrobial Mechanisms. Angew Chemie. 2017;129: 8211–8215. doi:10.1002/ange.201702313

26. Bennett RC, Oh MW, Kuo SH, Belo Y, Maron B, Malach E, et al. Random Peptide Mixtures as Safe and Effective Antimicrobials against Pseudomonas aeruginosa and MRSA in Mouse Models of Bacteremia and Pneumonia. ACS Infect Dis. 2021;7: 672–680. doi:10.1021/acsinfecdis.0c00871

27. Bishop BM, Juba ML, Devine MC, Barksdale SM, Rodriguez CA, Chung MC, et al. Bioprospecting the American alligator (Alligator mississippiensis) host defense peptidome. PLoS One. 2015;10: 1–17. doi:10.1371/journal.pone.0117394

28. König E, Zhou M, Wang L, Chen T, Bininda-Emonds ORP, Shaw C. Antimicrobial peptides and alytesin are co-secreted from the venom of the Midwife toad, Alytes maurus (Alytidae, Anura): Implications for the evolution of frog skin defensive secretions. Toxicon. 2012;60: 967–981. doi:10.1016/j.toxicon.2012.06.015

29. Conlon JM, Sonnevend A, Davidson C, David Smith D, Nielsen PF. The ascaphins: A family of antimicrobial peptides from the skin secretions of the most primitive extant frog, Ascaphus truei. Biochem Biophys Res Commun. 2004;320: 170–175. doi:10.1016/j.bbrc.2004.05.141

30. Mandal SM, Dey S, Mandal M, Sarkar S, Maria-Neto S, Franco OL. Identification and structural insights of three novel antimicrobial peptides isolated from green coconut water. Peptides. 2009;30: 633–637. doi:10.1016/j.peptides.2008.12.001

31. Yokoyama S, Kato K, Koba A, Minami Y, Watanabe K, Yagi F. Purification, characterization, and sequencing of antimicrobial peptides, Cy-AMP1, Cy-AMP2, and Cy-AMP3, from the Cycad (Cycas revoluta) seeds. Peptides. 2008;29: 2110–2117. doi:10.1016/j.peptides.2008.08.007

32. Tareq FS, Lee MA, Lee HS, Lee JS, Lee YJ, Shin HJ. Gageostatins A-C, antimicrobial linear lipopeptides from a marine Bacillus subtilis. Mar Drugs. 2014;12: 871–885. doi:10.3390/md12020871

33. Ward H, Perron GG, MacLean RC. The cost of multiple drug resistance in Pseudomonas aeruginosa. J Evol Biol. 2009;22: 997–1003. doi:10.1111/j.1420-9101.2009.01712.x

34. P. Lb, Michael Z, Jens R. Antimicrobial peptides: Application informed by evolution. Science (80-). 2020;368: eaau5480. doi:10.1126/science.aau5480

35. Imamovic L, Sommer MOA. Use of collateral sensitivity networks to design drug cycling protocols that avoid resistance development. Sci Transl Med. 2013;5. doi:10.1126/scitranslmed.3006609

36. Christian M, K. Gh, Nilsson WAI, H. Wh, A. Smo. Prediction of resistance development against drug combinations by collateral responses to component drugs. Sci Transl Med. 2014;6: 262ra156–262ra156. doi:10.1126/scitranslmed.3009940

37. Nichol D, Rutter J, Bryant C, Hujer AM, Lek S, Adams MD, et al. Antibiotic collateral sensitivity is contingent on the repeatability of evolution. Nat Commun. 2019;10. doi:10.1038/s41467-018-08098-6

38. Shaw LN, Aish J, Davenport JE, Brown MC, Lithgow JK, Simmonite K, et al. Investigations into σB-modelated regulatory pathways governing extracellular virulence determinant production in Staphylococcus aureus. J Bacteriol. 2006;188: 6070–6080. doi:10.1128/JB.00551-06

39. Hayouka Z, Chakraborty S, Liu R, Boersma MD, Weisblum B, Gellman SH. Interplay among Subunit Identity, Subunit Proportion, Chain Length, and Stereochemistry in the Activity Profile of Sequence-Random Peptide Mixtures. J Am Chem Soc. 2013;135: 11748–11751. doi:10.1021/ja406231b

40. Topman-Rakover S, Malach E, Burdman S, Hayouka Z. Antibacterial lipo-random peptide mixtures exhibit high selectivity and synergistic interactions. Chem Commun. 2020;56: 12053–12056. doi:10.1039/d0cc04493h

41. Aubry-Damon H, Soussy CJ, Courvalin P. Characterization of mutations in the rpoB gene that confer rifampin resistance in Staphylococcus aureus. Antimicrob Agents Chemother. 1998;42: 2590–2594. doi:10.1128/aac.42.10.2590

