## Supporting information for "Combination treatment can hinder the evolution of resistance to antimicrobial peptides"

Table S1. Antimicrobial peptides sequences and antimicrobial activity against ancestral strain

| *AMP* | *Sequence* | *MIC (μg/ml)* |
|---|---|---|
| *Ovispirin* | KNLRRIIRKIIHIIKKYG | 50 |
| *Aurein 1.2* | GLFDIIKKIAESF | 50 |
| *Melittin* | GIGAVLKVLTTGLPALISWIKRKRQQ | 6.25 |
| *Pexiganan* | GIGKFLKKAKKFGKAFVKILKK | 12.5 |
| *Temporin A* | FLPLIGRVLSGIL | 6.25 |
| *Pardaxin* | GFFALIPKIISSPLFKTLLSAVGSALSSSGGQE | 100 |

Table S2. AMP interactions using checkboard assay

| *Combination* | *Average FICi* | *Min FICi* |
|---|---|---|
| *Pexiganan + Melittin* | indifference (1.18) | indifference (1) |
| *Temporin + Melittin* | indifference (1.78) | indifference (1) |
| *Temporin + Pexiganan* | indifference (1.03) | partial synergy (0.75) |

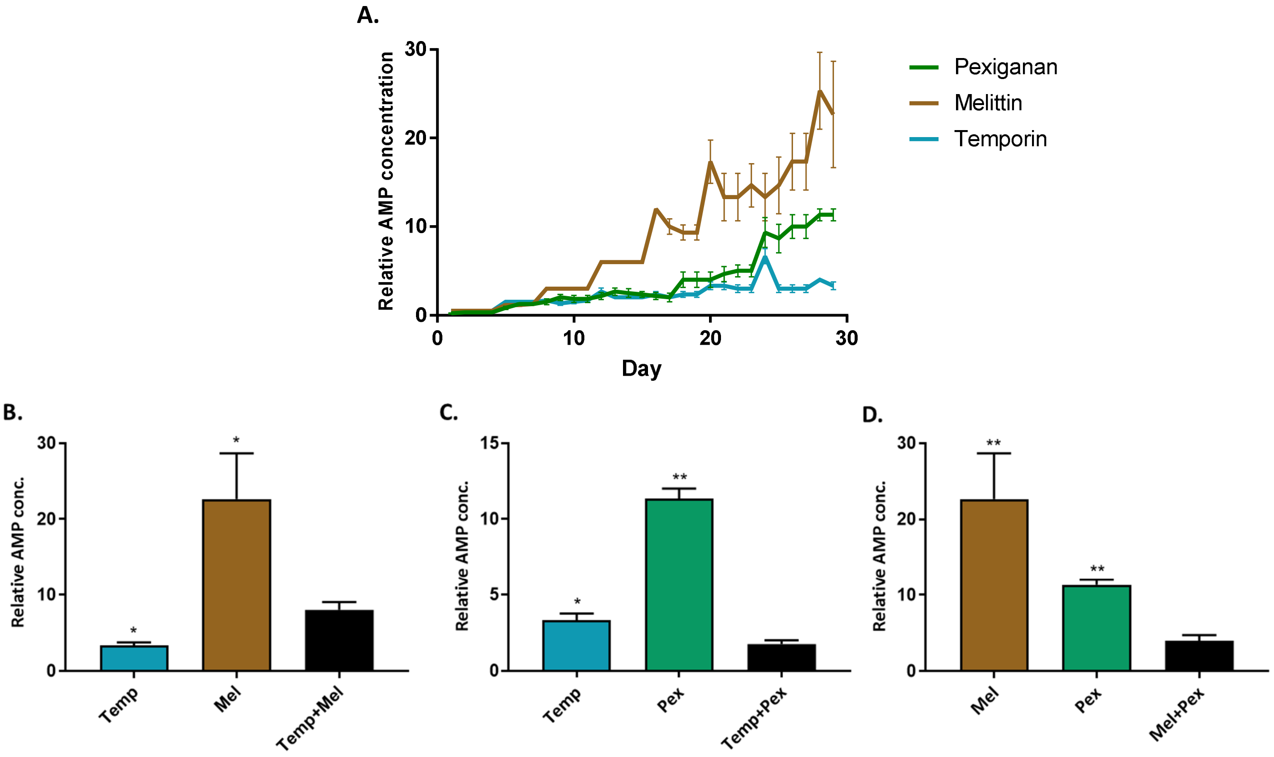


**Fig S1. Combination of AMPs can hinder the evolution of resistance.** Data represent the concentration of AMPs where bacteria grew after 29 transfers, normalized to initial MIC (n=6, Growth defined as $OD_{595}$>0.3). Each bar represents the mean of six lines + SEM. * p<0.05, ** p<0.01 (Mann-Whitney U test with Bonferroni correction, results compared to combinations strains) Mel, Melittin; Pex, Pexiganan; Temp, Temporin.

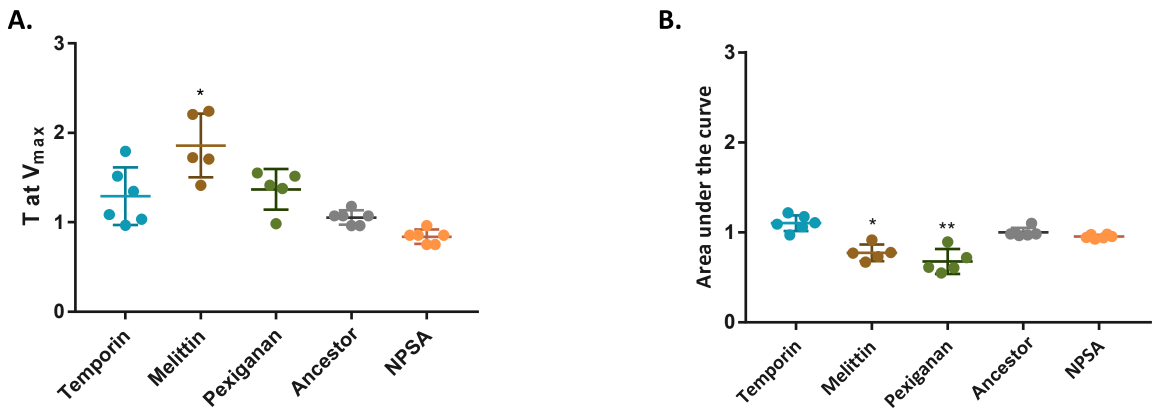


**Fig S2. Fitness cost of evolved strains**. Determination of fitness cost performed by growing the bacteria in the absence of AMPs. $OD_{595}$ was measured every 15 min through 24 hours. (A) Time to achieve maximal growth rate ($V_{max}$). (B) area under the curve. Values were calculated by plate reader software (Gen 5 and normalized to ancestor strain values). Each dot represents mean of triplicate, bars represent mean ± SD ($n_{melittin,\ pexiganan}$=5, Dunn's multiple comparison refed to ancestor strain, * p< 0.05, ** p< 0.01). NPSA- evolved strain without AMPs. The results represent three independent experiments.

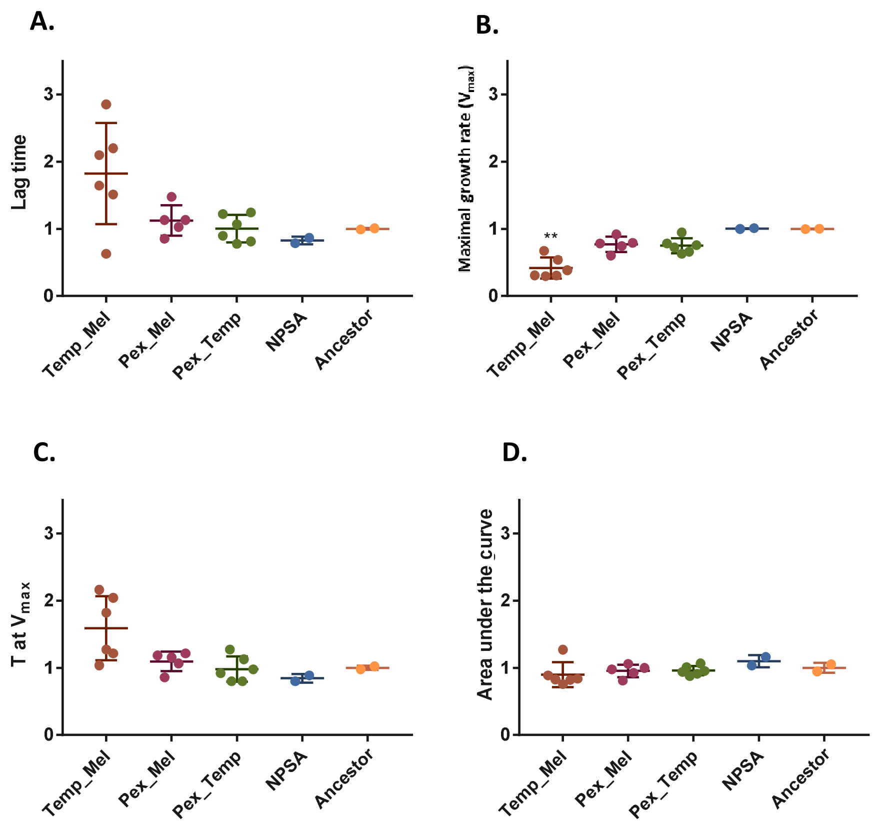


**Fig S3. Fitness cost of AMP-combinations evolved strains**. Determination of fitness cost performed by growing the bacteria in the absence of AMPs. $OD_{595}$ was measured every 15 min through 24 hours. (A) lag time. (B) maximal growth rate (Vmax). (C) time at Vmax. (D) area under the curves. A-C values were calculated by plate reader software (Gen 5 and normalized to ancestor strain values). Each dot represents mean of triplicate, bars represent mean ± SD (n=6, Dunn's multiple comparison refed to ancestor strain, ** p< 0.01). NPSA- evolved strain without AMPs. The results represent three independent experiments.

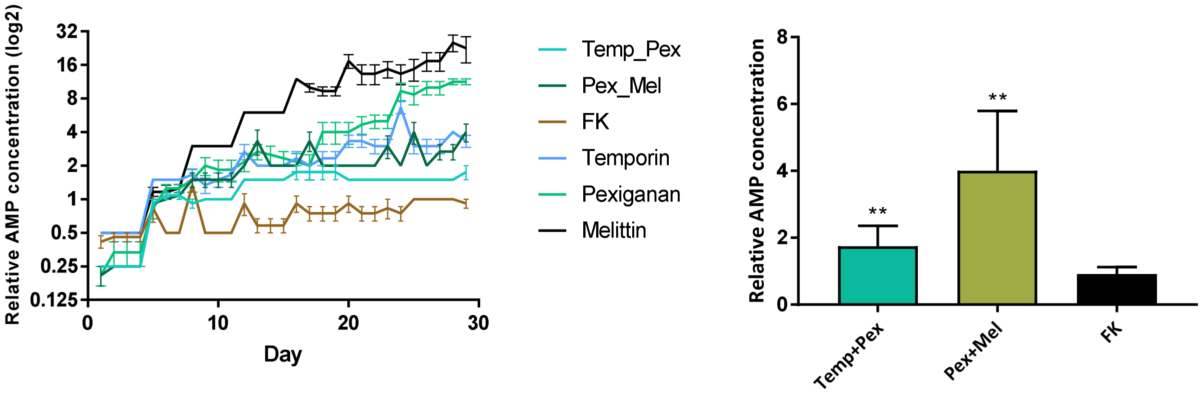


**Fig S4. Random peptides mixture composed of phenylalanine and lysine (FK) delays the evolution of resistance.** (A) Relative AMP concentration through the evolution (B) Comparison of the relative AMP concentration after 29 transfers, ** $p<0.01$ ($n=6$, Mann-Whitney U test with Bonferroni correction). Bars represent the mean + SEM.

Table S3. Statistical analysis

| *Test* | *n* | *Samples* | *Test value (U/ Z)* | *Adjusted p-values* | *Summary* |
|---|---|---|---|---|---|
| Kruskal-Wallis | 6 ($n_{mel}$=5) | All strains | 29.29 (Kruskal-Wallis) | <0.0001 | ** |
| Dunn's multiple comparison | 6,6 | Temp+Mel vs. Temp | 1.938 | 0.1052 | ns |
| | 6,6 | Temp+Mel vs. Mel | 1.772 | 0.1528 | ns |
| | 6,6 | Temp+Pex vs. Temp | 1.778 | 0.1509 | ns |
| | 6,6 | Temp+Pex vs. Pex | 3.889 | 0.0002 | ** |
| | 6,6 | Mel+Pex vs. Mel | 3.649 | 0.0005 | ** |
| | 6,6 | Mel+Pex vs. Pex | 2.277 | 0.0455 | * |
| MWU with Bonferroni correction | 6,6 | Mel vs. Pex+Mel | 0 | 0.0044 | ** |
| | 6,6 | Mel vs. Temp+Mel | 2 | 0.0174 | * |
| | 6,6 | Pex vs. Pex+Mel | 2.5 | 0.0216 | * |
| | 6,6 | Pex vs. Pex+Temp | 0.5 | 0.0086 | ** |
| | 6,6 | Temp vs. Mel+Temp | 7 | 0.1212 | ns |
| | 6,6 | Temp vs. Pex+Temp | 0 | 0.0044 | ** |
| Dunn's multiple comparison | 6,6 | Ancestor vs. Temporin | 1.507 | 0.6585 | ns |
| | 6,5 | Ancestor vs. Melittin | 3.162 | 0.0078 | ** |
| | 6,5 | Ancestor vs. Pexiganan | 1.835 | 0.3323 | ns |
| | 6,6 | Ancestor vs. NPSA | 1.189 | >0.9999 | ns |
| Dunn's multiple comparison | 6,6 | Ancestor vs. Temporin | 1.102 | >0.9999 | ns |
| | 6,5 | Ancestor vs. Melittin | 3.256 | 0.0057 | ** |
| | 6,5 | Ancestor vs. Pexiganan | 3.919 | 0.0004 | ** |
| | 6,6 | Ancestor vs. NPSA | 2.377 | 0.0873 | ns |
| Dunn's multiple comparison | 6,6 | Ancestor vs. Temporin | 1.233 | >0.9999 | ns |
| | 6,5 | Ancestor vs. Melittin | 3.04 | 0.0118 | * |
| | 6,5 | Ancestor vs. Pexiganan | 1.662 | 0.4823 | ns |
| | 6,6 | Ancestor vs. NPSA | 1.45 | 0.7347 | ns |
| Dunn's multiple comparison | 6,6 | Ancestor vs. Temporin | 0.9856 | >0.9999 | ns |
| | 6,5 | Ancestor vs. Melittin | 2.576 | 0.05 | * |
| | 6,5 | Ancestor vs. Pexiganan | 3.107 | 0.0095 | ** |
| | 6,6 | Ancestor vs. NPSA | 1.015 | >0.9999 | ns |
| MWU with Bonferroni correction | 6,6 | FK vs. Pex + Temp | 0 | 0.0044 | ** |

Statistical analysis performed using GraphPad Prism 7.02. * p<0.05, ** p<0.01
